# History-dependent muscle resistance to stretch remains high after small, posturally-relevant pre-movements

**DOI:** 10.1101/2022.12.22.521697

**Authors:** Brian C. Horslen, Gregory N. Milburn, Kyle P. Blum, Surabhi N. Simha, Kenneth S. Campbell, Lena H. Ting

## Abstract

The contributions of intrinsic muscle fiber resistance during mechanical perturbations to standing and other postural behaviors are unclear. Muscle stiffness, a traditional metric for estimating muscle’s intrinsic resistance to stretch, is known to vary depending on the current level and history of the muscle’s activation, as well as the muscle’s recent movement history; this property has been referred to as history dependence or muscle thixotropy. However, we currently lack sufficient data about the degree to which muscle stiffness is modulated across posturally-relevant characteristics of muscle stretch and activation. Here, we characterized the history dependence of muscle’s resistance to stretch in single, permeabilized, activated, muscle fibers in posturally-relevant stretch conditions and activation levels. We used a classic paired muscle stretch paradigm, varying the amplitude of a “conditioning” triangular stretch-shorten cycle followed by a “test” ramp-and-hold imposed after a variable inter-stretch interval. We tested low (<15%), intermediate (15-50%) and high (>50%) muscle fiber activation levels, evaluating short-range stiffness and total impulse in the test stretch. Muscle fiber resistance to stretch remained high at conditioning amplitudes of <1% L0 and inter-stretch intervals of >1 s, characteristic of healthy standing postural sway. A ~70% attenuation of muscle resistance to stretch was reached at conditioning amplitudes of >3% L0 and inter-stretch intervals of <0.1s, characteristic of larger, faster postural sway in balance-impaired individuals. Overall, amplitude and inter-stretch interval interact to disrupt myofilaments such that intrinsic resistance to stretch is attenuated if the stretch is large enough and/or frequent enough.

**Summary Statement:** Intrinsic muscle fiber resistance to stretch is preserved after small, slow pre-movements based on healthy postural sway, but markedly reduced as pre-movements increase to emulate abnormal postural sway.

## Introduction

Mechanical properties of contractile muscle, in concert with neural control mechanisms, are important mediators of postural control of the limbs and body (Nichols and Houk, 1976, 1973). When a muscle is stretched during standing balance due to mechanical perturbations to the body, it can take tens to thousands of milliseconds for the nervous system to initiate and complete a postural correction (Carpenter et al., 2005; Horak, 2006; Ivanenko and Gurfinkel, 2018; Ting et al., 2009). As a result of these relatively long sensorimotor delays, the initial stabilization of the body in response to a perturbation depends on resistive forces from intrinsic muscle mechanical properties (De Groote et al., 2017; Ivanenko and Gurfinkel, 2018; Nichols and Houk, 1973; Ting et al., 2009). Muscle stiffness, a traditional metric for estimating muscle’s intrinsic resistance to stretch, is known to vary depending on the current level and history of the muscle activation, as well as the muscle’s recent movement history (Campbell and Moss, 2002). However, we currently lack sufficient data about the degree to which muscle stiffness is modulated across posturally-relevant characteristics of muscle stretch and activation. Therefore, the goal of this study was to characterize muscle’s intrinsic resistance to stretch in single muscle fiber experiments at muscle stretch and activation levels relevant to standing postural sway in humans in health and disease.

History dependence of muscle stiffness due to muscle stretch has been well-characterized in single muscle fiber experiments. When a muscle fiber is stretched, it exhibits a high but transient initial stiffness (stress increase per unit change in muscle fiber length) referred to as short-range stiffness (SRS). If the muscle continues to be stretched beyond this “short range”, then the stiffness decreases (Rack and Westbury, 1974). History-dependant variations in this short-range stiffness i.e., muscle thixotropy has been demonstrated in single muscle fibers using a paired stretch paradigm where a conditioning stretch-shorten cycle immediately precedes a test stretch (Fig. 1A) (Herbst, 1976; Lakie and Campbell, 2019; Lakie and Robson, 1988; Lännergren, 1971). A conditioning stretch-shorten cycle of sufficient amplitude (often ~3% optimal fiber length, L0) reduces SRS in the second, test stretch, of the same amplitude by up to ~50% (Campbell and Moss, 2002; Lännergren, 1971). If the inter-stretch interval between the conditioning and test stretch is increased, the SRS in the test stretch increases monotonically up to at least 10 s before fully recovering its stiffness (Campbell and Moss, 2002; Rassier and Herzog, 2004).

**Figure 1:**
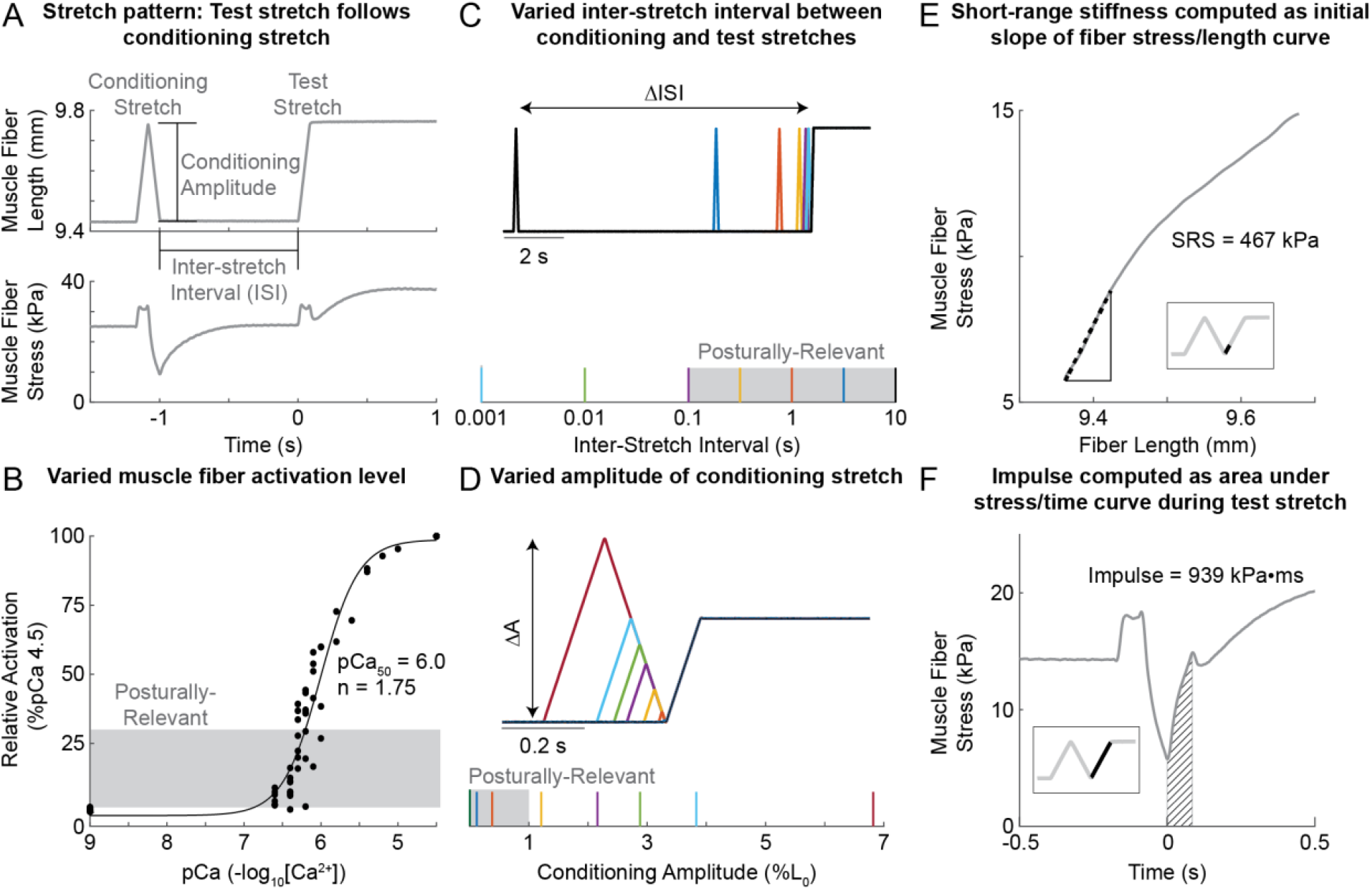
Experimental design: muscle fiber stretch patterns, activation, and outcome measures. A) Exemplar data illustrating the muscle stretch response during the paired conditioning-test stretch protocol for a fiber activated to 12.5% max in pCa 6.4 solution. Muscle fiber length was changed to impose a “conditioning” stretch-shorten cycle of variable amplitude followed by a ramp and hold “test” stretch, with a variable inter-stretch interval. Modulation of muscle resistance was characterized in the test stretch. B) Muscle fiber activation level versus Ca^2+^ concentration of activating solution for all fibers included in the data set. Relative activation was calculated as a percentage of isometric force produced in maximally activating pCa 4.5 solution. These data were used to identify [Ca^2+^] that yielded posturally-relevant activation levels (2-30% max), shaded in gray. Single-trial samples from all fibers (black dots) were used to identify Hill equation (Equation 1) parameters pCa_50_ = 6.00 and n = 1.75, representing mean activation characteristics across all fibers in our sample. C) Variations in inter-stretch interval, i.e., the duration of the isometric hold between the conditioning stretch-shorten cycle and ramp-and-hold test stretch. Posturally-relevant ISI shaded in gray at the bottom. D) Variations in conditioning stretch-shorten cycle amplitude with constant stretch velocity. Posturally-relevant amplitudes shaded in gray at the bottom. E) Short-range stiffness (SRS) was calculated as the slope of the muscle fiber stress-strain curve over the first 20 ms of stretch (black dashed line; see inset for corresponding region of length-time trace). F) Impulse was calculated as the area under the stress-time curve (dashed region) for the duration of the ramp period (see inset for corresponding region of length-time trace).

Much of the knowledge on muscle thixotropy comes from experiments conducted at high levels of muscle activation rather than on lower, posturally-relevant levels. During quiet standing, plantar flexor muscles are typically activated between 5% and 20% of maximum voluntary activation (Warnica et al., 2014) in healthy and impaired adults, respectively. While larger history-dependant reductions in muscle SRS have been demonstrated at intermediate activation levels (~20 to 50% maximum activation) (Campbell and Moss, 2002), most muscle fiber thixotropy studies focus on 50-100% activation levels so as to maximize signal-to-noise ratio, or suit other experimental needs (Altman et al., 2015; Lynch et al., 2008). Therefore, in the current study we focused on the effects of activation on muscle thixotropy when activated below 50%.

Prior experiments on muscle thixotropy have not investigated the effect of conditioning stretch amplitude at small, posturally-relevant amplitudes. In healthy standing balance, the ankle dorsi- and plantar flexors are critical for maintaining upright posture and are continuously stretched and shortened as the body sways (Day et al., 2013; Loram et al., 2009). Postural sway-induced ankle muscle stretch during unperturbed standing is estimated to be less than 1% L0 in healthy adults (Day et al., 2013; Ivanenko and Gurfinkel, 2018; Loram et al., 2009). However, postural sway is faster and larger in individuals with impaired balance (Horak and Macpherson, 2011; Maurer et al., 2003); increases in postural sway of up to two times that of healthy individuals (Maurer et al., 2003) suggest that muscle stretch could reach up to 2% in individuals with balance (Loram et al., 2009). This study of conditioning amplitudes between 1-3% L0 may thus be critical to understanding how muscle mechanical contributions to balance control differ across healthy and impaired conditions.

Based on the biophysical mechanisms underlying muscle force generation, we predicted that muscle resistance to stretch would be modulated by posturally-relevant timing and amplitudes of the conditioning stretch at low activations typical of standing balance control (Lakie and Campbell, 2019). Briefly, muscle force can be considered to arise based on the stiffness and length of bound actin-myosin muscle cross-bridges. The greater the number of attached cross-bridges, the greater the muscle stiffness. Muscle force increases based on the length of the stretched cross-bridges. Muscle short-range stiffness arises when the number of bound cross-bridges is high, such as when the muscle is held isometrically; muscle stiffness rapidly decreases when sarcomeres are stretched beyond a few nanometers, causing myosin heads to be pulled off their actin binding sites, rapidly reducing the number of bound cross bridges. As such, stretching a muscle decreases the number and length of bound cross bridges, reducing muscle stiffness and force in subsequent stretches. Conversely, increasing the inter-stretch interval allows cross-bridges to reattach over time, increasing muscle resistance to stretch in subsequent stretches. Therefore, the muscle resistance to stretch will depend on the net effect of prior stretch and rest intervals.

While studies on the mechanisms of history-dependence in muscle resistance to stretch focus on short-range stiffness, the muscle’s force throughout a stretch provides more information about its role in movement. Behaviourally, the success of a muscle in rejecting a postural perturbation depends on its force throughout stretch. Impulse, measured as the time integral of force, is proportional to the change in momentum imparted to an inertial load, and may provide better insight into the role of history-dependent muscle resistance to stretch on behavior.

Here we characterized the history-dependence of single muscle fiber resistance to stretch in posturally-relevant ranges of inter-stretch interval, muscle activation, and conditioning stretch amplitude. We activated permeabilized muscle fibers and used a paired conditioning-test stretch design where a balance perturbation-like “test” stretch was preceded by a postural sway-like stretch-shorten “conditioning” cycle (Fig. 1A). We systematically varied 1) bath Ca^2+^ concentration to alter muscle activation level (Fig. 1B), 2) the inter-stretch interval between conditioning stretch-shorten cycles and the test stretch (Fig. 1C), and 3) the amplitude of conditioning stretches (Fig. 1D). We used a constant velocity for all stretches. We characterized muscle resistance to stretch in two ways. We estimated 1) short-range stiffness at the onset of stretch (Fig. 1E), and 2) overall resistance of the muscle to stretch, calculated as total impulse throughout the duration of the test stretch (Fig. 1F). Overall, we found that muscle resistance to stretch is highest at conditions commensurate with healthy postural muscle activation and sway but reduced most by prior movement at higher postural activation and sway levels characteristic of balance impairments.

## Methods

### Ethical Approval

Eleven soleus muscle fibers harvested from 2 adult female Sprague-Dawley rats (Envigo, Indianapolis, IN, USA; RRID: RGD_737903) were studied. Experiments were performed at The University of Kentucky. Animals were housed in clean cages with a 12hr light/dark cycle in an Animal Care Facility and had access to food and water ad libitum. Experimental procedures were reviewed and approved by the University of Kentucky IACUC (#: 00784M2004; Assurance number D16-00217 [A3336-01]).

### Sample harvest and preparation

Procedures and solutions used for muscle fiber harvest, permeabilization, and storage, as well as equipment and procedures used to conduct experiments are described in detail elsewhere (Campbell and Moss, 2002; Campbell, 2006). In brief, soleus muscles were harvested immediately after sacrificing the animal, teased into fiber bundles, then permeabilized for 4 hours. Permeabilized muscle fiber bundles were stored in a glycerol solution at −20°C for no more than 1 month prior to use.

### Experimental apparatus

Single muscle fibers were tied between a motor (model 312, Aurora Scientific Inc., Aurora, ON, Canada) and force transducer (model 403, Aurora Scientific Inc.). Muscle fiber length was controlled, and length and force data recorded, using SLControl software (www.slcontrol.com; (Campbell, 2006)); both data sampling and length position update rates were set to 1kHz. Initially, muscle fiber length was manually adjusted under microscope and in minimally activating solution (see below) to a mean sarcomere length of 2.6μm (assumed optimal fiber length; L_0_). This fiber length was recorded and all subsequent length manipulations were scaled as a fraction of this L_0_.

### Characterizing muscle fiber activation

After establishing L_0_, we incrementally increased each muscle fiber bath Ca^2+^ concentration to characterize the fiber’s full range of activation levels (Fig. 1B). Permeabilized fibers were chemically activated using free Ca^2+^ solutions with concentrations ranging from pCa (=-log_10_[Ca^2+^]) 9.0 (minimal Ca^2+^) to pCa 4.5 (maximal), with densest sampling in the pCa 6.6-6.0 range (Fig. 1B). pCa solutions contained 20 mM imidazole, 14.5 mM creatine phosphate, 7 mM EGTA, 4 mM MgATP, 1mM free Mg2+, free Ca^2+^ ranging from 1 nM to 32 μM and KCl to adjust ionic strength to 180 mM with pH 7.0 at 22° C; a 22° C mean temperature was maintained for all experiments. At each activation level we calculated mean isometric stress over 0.9 s with the fiber held at a mean sarcomere length of 2.6 μm (L_0_). We then calculated the stress as a percentage of pCa 4.5 isometric stress and took this value as the achieved activation level. We fit data for all fibers to a sigmoidal curve to estimate the mean Hill Equation parameters, *n* and pCa_50_ (pCa required to achieve 50% activation), for our sample (Fig. 1B) (Eqn. 1) (Donaldson and Kerrick, 1975). We estimated that pCa_50_ = 6.00 and n = 1.75.

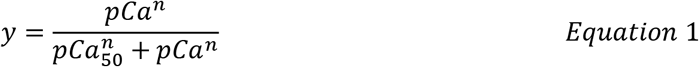

### Identifying posturally-relevant activation levels

After characterizing fiber Ca^2+^ sensitivity, we selected at least three pCa solutions to test the effects of muscle fiber activation level on history-dependant muscle resistance to stretch. pCa solutions were selected to yield low, intermediate, and high activation levels relative to muscle activation in standing balance tasks (2-30%) (Warnica et al., 2014). Low activation was defined as less than 15% max (where max = isometric force at pCa 4.5); intermediate was 15-40% max; and high activation was greater than 40% max. To preserve the viability of the muscle fibers, activation levels were tested in ascending order, where all conditioning amplitude and inter-stretch interval combinations were tested (c.f. *Posturally-relevant inter-stretch intervals* and *Posturally-relevant conditioning amplitudes*) before moving to the next activation level. Activation levels were verified post-hoc and used to assign groups for statistical comparison. Not all fibers yielded data in each activation range and some fibers yielded more than one data set within a range. If fiber appearance or mechanical properties had not deteriorated after stretching at high activation (i.e., striations remained clear under the microscope and isometric force was not changed by more than 20% from initial activation) (Campbell and Moss, 2002), then the fibre was maximally activated in pCa 4.5 solution. At pCa 4.5, the muscle was stretched for as many trials as possible before degradation (usually 2-3 stretches) to create a maximal activation data set.

### Paired conditioning and test stretch protocol

To characterize the modulation of muscle resistance to stretch, inter-stretch interval and conditioning stretch-shorten amplitude were systematically varied (Fig. 1C,D, c.f. *Posturally-relevant inter-stretch intervals* and *Posturally-relevant conditioning amplitudes*), while analyses were performed on the test stretch (Fig. 1E,F). Ramp and hold characteristics of the test stretch were fixed across all trials (amplitude: 3.83% L_0_, velocity: 45.45 % L_0_/s, hold duration: 2 s); these parameters were selected for behavioral relevance to postural perturbations. We tested 25 unique inter-stretch interval and conditioning amplitude combinations in 11 fibres. This included a test stretch with no preceding conditioning stretch to establish a baseline response at each activation level. In 3 of the 11 fibers, we also included an additional 25 combinations to further explore interactions between inter-stretch interval and conditioning amplitude (c.f. *Posturally-relevant inter-stretch intervals* and *Posturally-relevant conditioning amplitudes*).

### Posturally-relevant inter-stretch intervals

Inter-stretch interval (ISI) between the conditioning and test stretches were varied along a logarithmic scale between 0.001 s and 10 s (Fig. 1A, C). This range of inter-stretch intervals (0.001 s, 0.1 s, 0.316 s, 1 s, 3.162 s, 10 s) was selected considering known ISI-dependent changes in muscle stiffness (Campbell and Moss, 2002, 2000; Proske and Stuart, 1985) and the range of human postural sway frequencies (0.1-10 Hz; Fib. 1C gray shading) (Prieto et al., 1996). In a sub-set of fibers (3/11), we included a sample at a 0.01 s inter-stretch interval to increase the resolution of time-dependent effects.

### Posturally-relevant conditioning amplitudes

We systematically varied the conditioning stretch amplitude in the paired stretch protocol (Fig. 1A, D). Conditioning stretch velocity was fixed at 45.45% L_0_/s, matching the test stretch velocity. Based on pilot studies, we anticipated a logarithmic scaling of responses with conditioning stretch amplitude, causing us to sample a range from 0.12% L_0_ to 3.83% L_0_ (Fig. 1D; 0.12%, 0.38%, 1.21%, 3.83% L_0_). This range encompasses very small stretches that are less than the short-range elastic limit (0.2% L_0_) (Hill, 1968), small amplitudes observed in postural sway (<1% L_0_; Fig. 1D gray shading), amplitudes used in previous muscle fiber mechanics studies (3% L_0_) (Campbell and Moss, 2002), and large, dynamic, amplitudes that occur in reaching or gait (> 3% L_0_). In a sub-set of fibers (3/11), we added trials with conditioning amplitudes of 2.16%, 2.88% and 6.83% L_0_ to improve resolution and to better establish the interacting effects of conditioning stretch interval and amplitude.

### Muscle resistance outcome measures

We quantified history-dependant changes in muscle fiber resistance to stretch by calculating short-range stiffness (SRS) and impulse of the test stretch in each trial. SRS refers to the initial stiffness of the muscle in response to stretch and was calculated as the slope of the stress vs. strain curve in the first 0.02 s after the onset of test stretch (Fig. 1E). Impulse refers to the time-integral of force and characterizes the total resistance of the muscle throughout the test stretch, well beyond the initial period where SRS is computed. Impulse was computed as the area under the stress vs. time curve for the duration of the lengthening phase of the ramp stretch (0.084 s) (Fig. 1F).

### Statistical analyses

We performed an initial assessment of the effects of muscle fiber activation level on history-dependant changes in SRS and impulse. SRS and impulse were computed for unconditioned test stretches across activation levels, as well as test stretches conditioned with a 3.83 % L_0_, 0.001s ISI at all activation levels. Such large-amplitude, short interval conditioning patterns have previously been shown to yield large reductions in SRS across a range of fiber activation levels (Campbell and Moss, 2002). Separate linear mixed models were used to test whether SRS or impulse varied with activation and conditioning stretch. In each model, activation was treated as a continuous variable, conditioning was a dichotomous variable (conditioned versus unconditioned), and each fiber (j) was included as a random variable.

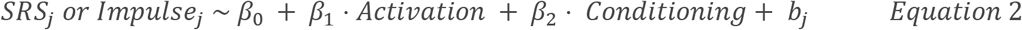

We first tested the effects of activation on absolute levels of SRS and impulse and compared these effects in conditioned versus unconditioned test stretches. We further tested the effect of activation on relative changes in SRS (% unconditioned test stretch) and impulse. Here we used categorical levels of fiber activation of low, intermediate, high, and max as independent variables, and the relative change in conditioned versus unconditioned SRS and impulse (% Unconditioned) as dependent variables in the linear mixed model. In all our analyses we report the coefficient values, t-values (estimate/standard error) and p-values. Analyses were performed in R software using “lmerTest::lmer” routines.

We next tested how inter-stretch interval, conditioning amplitude, and muscle fiber activation level affect history-dependant changes in relative SRS or impulse (Fig 4). We computed SRS and impulse for all test stretches, and expressed them as % unconditioned test stretch, to obtain relative SRS and impulse values. On the full data set (n=11 fibers), we used linear mixed models in R software (lmerTest::lmer) with inter-stretch interval (6 levels: 0.001 s, 0.1 s, 0.316 s, 1 s, 3.162 s, 10 s), conditioning amplitude (4 levels: 0.12 % L_0_, 0.38 % L_0_, 1.21 % L_0_, 3.83 % L_0_) and activation level (3 levels: low, intermediate, high) coded as categorical variables, fiber as a random variable, and the dependent measures, SRS or impulse as continuous variables.

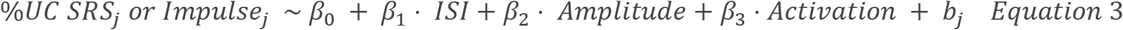

**Figure 2:**
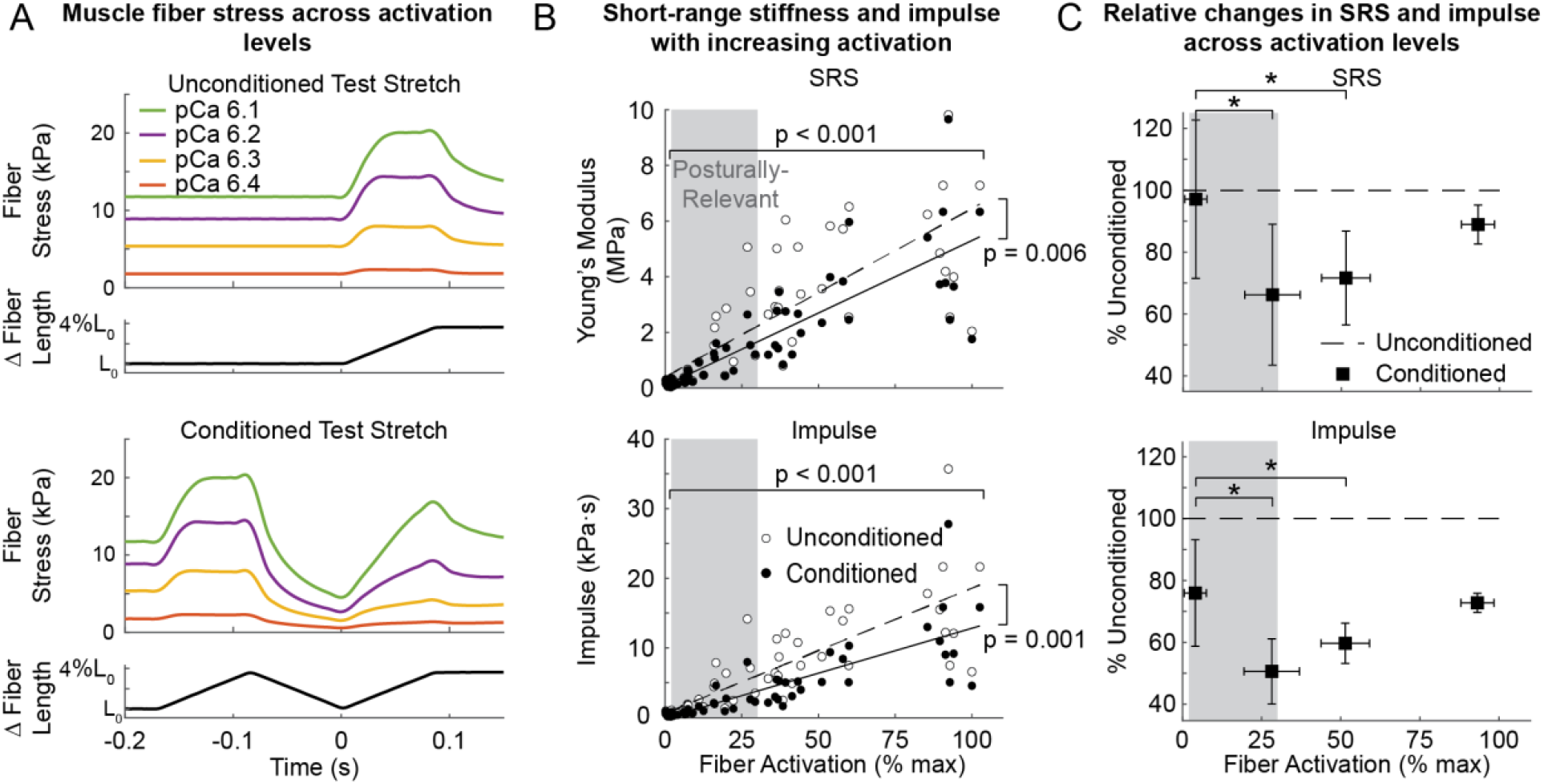
Effect of activation level on thixotropic changes to muscle fiber resistance to stretch. A) Representative data from a single permeabilized muscle fiber demonstrate that muscle fiber stress increased with activation level (compare colors) in both unconditioned (top) and conditioned trials (bottom) with a 3.83% L_0_ conditioning stretch and 0.001 s inter-stretch interval. B) Across all fibers, short-range stiffness, SRS, (top) and impulse (bottom) increased monotonically with increasing muscle fiber activation (slope of line: p<0.001) for both unconditioned (open circles) and conditioned stretches (black dots). The SRS of unconditioned stretches increased faster than that of conditioned stretches (slope of line: p=0.005). C) The largest relative decreases in both SRS (top) and impulse (bottom) between conditioned (black squares) and unconditioned trials (dashed horizontal line at 100%) occurred at intermediate activation levels. Error bars represent SD and horizontal bars with asterisks represent significant differences (p<0.05) between activation levels; gray boxes represent posturally-relevant activations ranges (2-30% max)

**Figure 3:**
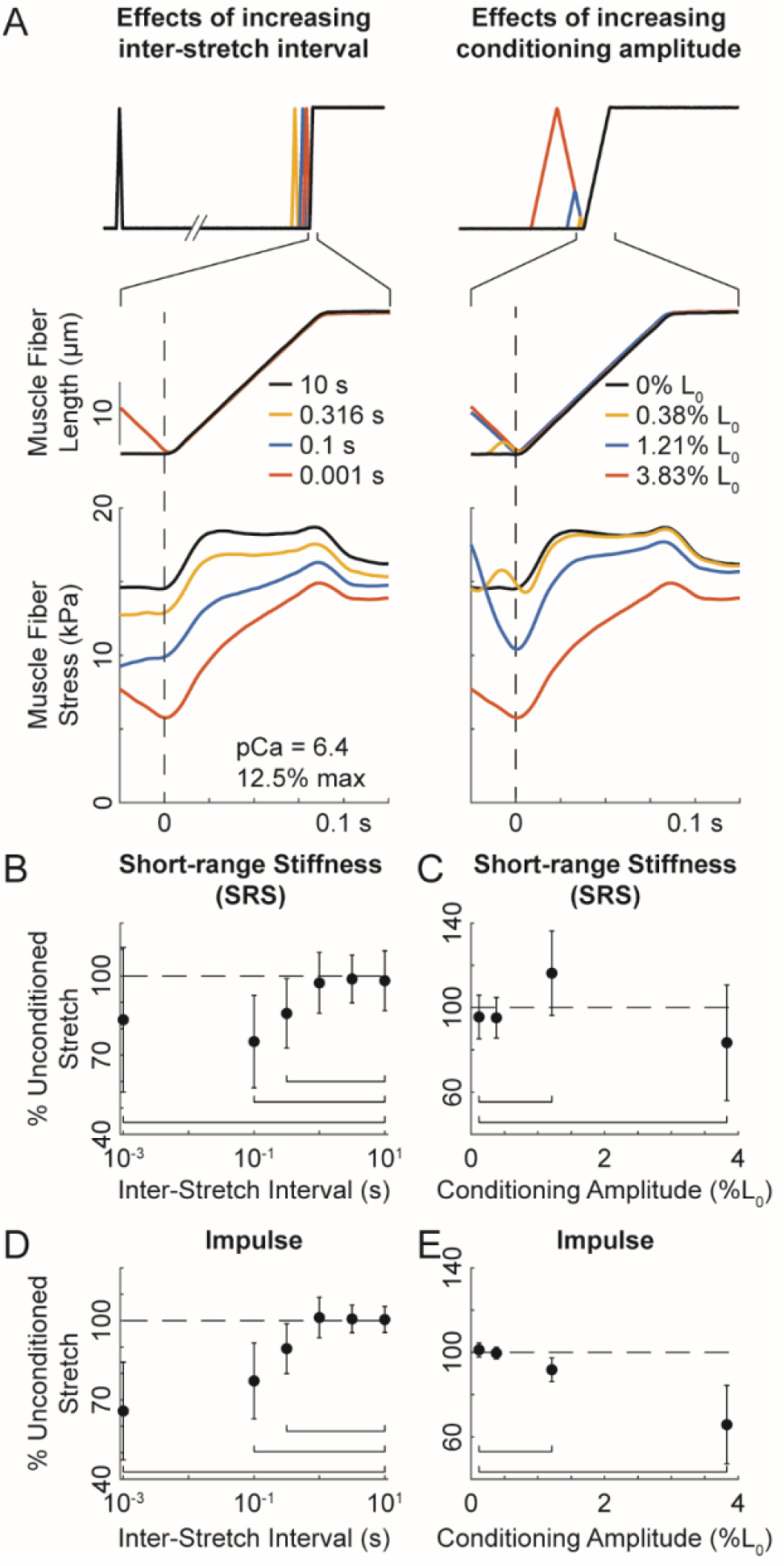
Independent effects of inter-stretch interval and conditioning stretch amplitude on muscle fiber stiffness. A) Representative data of muscle fiber stress at low activation (12.5%, pCa 6.4) showing the independent effects of decreasing inter-stretch interval when conditioning amplitude is constant (3.83% L_0_; left column, top panel), and increasing conditioning stretch amplitude when inter-stretch interval is constant (0.001 s; right column, top panel). Traces are aligned in time to onset of the test stretch (dashed vertical lines), which was identical in velocity and displacements across all trials. The SRS in conditioned test stretches was greatest when inter-stretch interval was longest (10 s; left column, black traces), and decreased with shorter ISIs (compare colors). Conversely, SRS was greatest when the conditioning amplitude was smallest (0% L_0_; right column, black trace), and decreased as conditioning amplitude increased (compare colors). Across all fibers, the % unconditioned short-range stiffness (SRS) was lowest at B) short inter-stretch intervals and C) large conditioning stretch amplitudes. Horizontal brackets indicate statistically significant (p<0.05) contrasts between sample and 10 s inter-stretch interval value. Likewise, % unconditioned impulse was lowest when D) inter-stretch interval was short or E) conditioning amplitude was large. Horizontal brackets indicate statistically significant (p<0.05) contrasts between sample and 0.12% L_0_ conditioning amplitude value. Error bars indicate SD.

**Figure 4:**
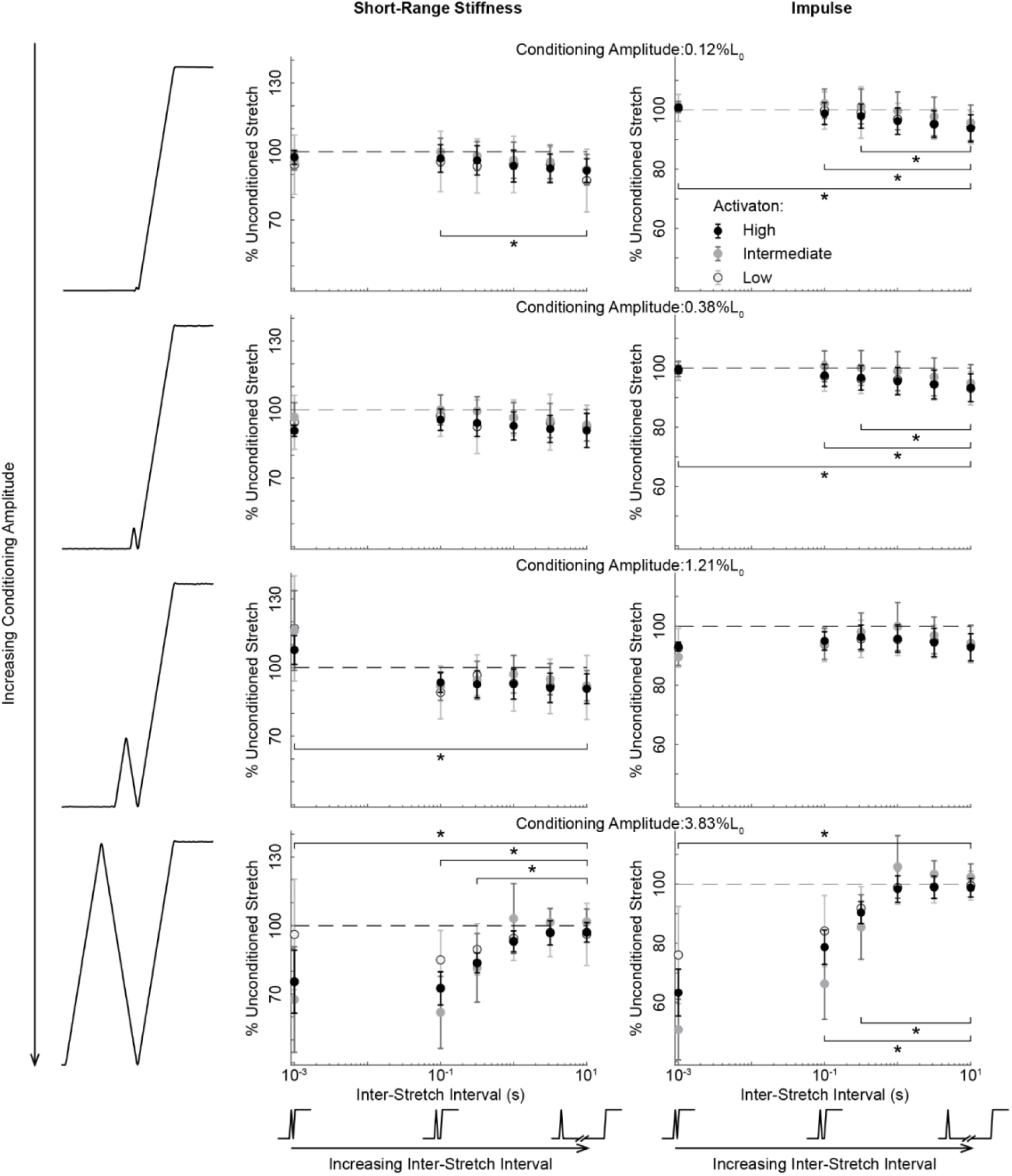
Interactions between conditioning stretch amplitude and inter-stretch interval across activation level. Each panel shows the effects of increased inter-stretch interval at three activation levels for a single conditioning stretch amplitude on mean short-range stiffness (SRS; left column) and impulse (right column). From top to bottom, each row represents progressively larger conditioning stretch amplitudes. At small conditioning stretch amplitudes (top three rows), there is little absolute change in either short-range stiffness or impulse with increasing inter-stretch interval. As conditioning stretch amplitude increases, both short-range stiffness and impulse are reduced with at the smallest inter-stretch intervals (bottom row). Horizontal dashed line indicated 100% unconditioned stretch SRS or impulse values. Error bars indicate standard deviation. Horizontal brackets and asterixis indicate statistically significant differences from the 10s inter-stretch interval condition, collapsed across activation conditions.

The maximum activation level did not yield enough ISI and conditioning amplitude combinations to be included in this analysis. The reference level was set to be the trials with the smallest conditioning amplitude (0.12% L_0_), longest ISI (10 s), and lowest activation level (<15% max).

We used post hoc contrasts to test our specific hypotheses about the effects of inter-stretch interval and conditioning amplitude on history-dependant changes in SRS and impulse (Fig 3A, bottom row). Estimated marginal means contrast tests (R software, emmeans::emmeans) were used to isolate main effects from the linear mixed model results as outlined in the above paragraph.

To isolate the effect of inter-stretch interval, we limited the analysis to the large, 3.83% L_0_, amplitude because small conditioning stretches often failed to change SRS or impulse (Fig 4) (Lännergren, 1971). We collapsed the data across all activation levels and fixed conditioning amplitude at 3.83% L_0_ and then compared change in SRS or impulse (% unconditioned) marginal means across inter-stretch intervals. To isolate the effect of conditioning amplitude on SRS or impulse, we again collapsed the data across activation levels and fixed ISI at 0.001 s, then compared SRS or impulse (% unconditioned) marginal means across amplitudes. Similar to ISI above, we limited the analysis to the short, 0.001 s, ISI because long inter-stretch intervals have been shown to fail to change SRS (Campbell and Moss, 2002). The threshold for statistical significance was set to 0.05; p values were adjusted using Tukey’s method for multiple comparisons (ISI: 6 estimates; conditioning amplitude: 4 estimates).

We performed similar analysis on the set of samples with higher sample density (n = 3) to further characterize ISI and conditioning amplitude interaction effects. As above, we used separate linear mixed models (lmerTest::lmer) to test for effects of activation and conditioning parameters on SRS and impulse. In these tests, we included categorical variables: inter-stretch interval (7 levels: 0.001 s, 0.01 s, 0.1 s, 0.316 s, 1 s, 3.162 s, 10 s); conditioning amplitude (7 levels: 0.12% L_0_, 0.38% L_0_, 1.21% L_0_, 2.16% L_0_, 2.88% L_0_, 3.83% L_0_, 6.82% L_0_); and activation level (3 levels: low, intermediate, high), and muscle fiber as a random variable. Again, the reference level was set to be the trials with the smallest conditioning amplitude (0.12% L_0_), longest ISI (10 s), and lowest activation level (<15% max).

## Results

### Effect of activation on the history dependence of muscle fiber resistance to stretch

Consistent with previous studies (Campbell and Moss, 2002), muscle resistance to stretch increased with muscle activation level in both unconditioned and conditioned test stretches and was lower in conditioned trials. The change in muscle stress over time during the test stretch changed in amplitude as bath calcium concentration increased (see example in Fig. 2A, colored traces) in both unconditioned and conditioned test stretches. However, there was a qualitative change in the shape of the muscle stress in response to the test stretch in conditioned (Fig. 2A, lower), compared to unconditioned (Fig. 2A, upper) responses. These qualitative changes reflect a history-dependent change in muscle resistance to stretch. Across all fibers in our sample both short-range stiffness and impulse increased with activation (Fig. 2B; short-range stiffness: β = 57.7, t_107.3_ = 17.9, p < 0.001; impulse: β = 159.3, t_107.3_=17.6, p < 0.001). However, both short-range stiffness and impulse were significantly greater in unconditioned stretches than conditioned ones when considered across all activations (SRS: β = 557.3, t_105_=2.8, p = 0.006; impulse: β = 2210, t_105_=3.9, p < 0.001).

Relative reduction in muscle resistance to stretch in the conditioned trials was nonlinear as a function of activation level, with the greatest reductions at posturally-relevant intermediate activation levels (Fig. 2C, gray shaded region). When activation was binned into low (<15% max), intermediate (15-40% max), high (40-60% max), and maximal activation levels (>60% max), the conditioned SRS and impulse expressed as a % unconditioned level exhibited a U-shaped relationship to activation (Fig. 2C). Short-range stiffness was decreased from unconditioned levels in the low activation bin (β = 96.5, t_23.9_ = 20.7, p < 0.001) and was further decreased (Fig. 2C upper) in the intermediate (β = −29.2, t_46.8_ = −4.4, p < 0.001) and high (β = −23.3, t_49.3_ = −2.8, p = 0.008) activation bins, compared to low activation; maximal activation short-range stiffness was not different from low activation values, when expressed as a percent of unconditioned values. Similarly, impulse was decreased in the low activation condition, compared to unconditioned levels (β = 75.9; t_30.5_ = 29.0; p < 0.001), and further decreased (Fig. 2C lower) in both the intermediate (β = −25.8; t_47.59_ = −6.1; p < 0.001) and high activation (β = −15.8; t_48.1_ = −3.0; p = 0.004) bins. Impulse, as a percentage of unconditioned stretch values, was not different from low activation in the maximal activation bin.

### Independent effects of inter-stretch interval and conditioning amplitude on muscle resistance to stretch

We observed similar qualitative changes in muscle stress over time during the test stretch due to independent variations in inter-stretch interval and conditioning amplitude (see example in Fig. 3A of a fiber at 12.5% activation). As shown previously, when conditioning amplitude was held constant at a large amplitude (3.83% L_0_) muscle stress was greatest at the longest inter-stretch interval of 10 s (Fig. 3A, black traces left panel). Muscle stress exhibited a similar profile when there was no conditioning stretch (0% L_0_; Fig. 3A right panel black trace). Muscle stress during the test stretch decreased systematically when either inter-stretch interval was decreased independently (Fig 3A, left column, colored traces), or conditioning stretch amplitude increased independently (Fig. 3A, right column, colored traces). In both cases, stress decreased both at the onset of the test stretch (dashed vertical line) as well as for the duration of the lengthening period. Note that the drawn conditions where the stress was lowest represent the same trial, where the inter-stretch interval was 0.001 s and conditioning stretch amplitude was 3.83% L_0_ (Fig. 3A red traces in left and right panels).

Across all fibers and activation levels, significant reductions in muscle resistance to stretch were found when decreasing inter-stretch interval while holding conditioning stretch amplitude constant. The reduction in % unconditioned short-range stiffness and impulse were both lower in the shortest three inter-stretch intervals (0.001 s, 0.01 s and 0.316 s) compared to the longest (10 s) inter-stretch interval (Fig. 3B, Fig. 3D). The largest reductions in both short-range stiffness and impulse were at the shortest inter-stretch interval where they were both around 70% unconditioned, and increased with longer inter-stretch intervals until reaching about 100% unconditioned at 0.316 s (Fig. 3B, Fig. 3D).

Across all fibers and activation levels, significant reductions in muscle resistance to stretch were also found when increasing conditioning stretch amplitude while holding inter-stretch interval constant. Large conditioning stretch amplitudes (3.83% L_0_) led to significant reductions in short-range stiffness (84% unconditioned) and impulse (66% unconditioned), compared to the smallest conditioning amplitude (0.12% L_0_; Fig. 3C). However, at the next smallest conditioning stretch amplitude of 1.21% L_0_ conditioning stretch amplitude, short-range stiffness was slightly higher than unconditioned levels, while impulse was slightly lower than unconditioned levels; data from smaller conditioning stretch amplitudes were about 100% unconditioned.

Interactions between the effects of inter-stretch interval and conditioning stretch amplitude were found at all activation levels. When all conditioning and activation permutations were examined, there were main effects of both inter-stretch interval (short-range stiffness: F_5,1283.8_ = 9.8 p < 0.001; impulse: F_5,1283.8_ = 42.8, p < 0.001) and conditioning stretch amplitude (short-range stiffness: F_3,1283.8_ = 20.0, p < 0.001; impulse F_3,1283.8_ = 109.0, p < 0.001), consistent with the main effects reported above. There was an interaction between inter-stretch interval and conditioning amplitude on short-range stiffness (F_15,1283.8_ = 21.8, p < 0.001) and impulse (F_15,1283.8_ = 72.0, p < 0.001). These interactions are visualized by plotting all changes in short-range stiffness and impulse as a function of inter-stretch interval at each of four conditioning stretch amplitudes (Fig. 4). The largest reductions in short-range stiffness and impulse are observed when the conditioning stretch amplitude was largest (Fig. 4, bottom row). There were also small, but statistically significant, increases in both short-range stiffness and impulse at small conditioning amplitudes. While we found an effect of activation level on short-range stiffness and impulse at the largest-amplitude, shortest-interval, stretches (Fig. 2C), there was no effect of activation on short-range stiffness or impulse when all inter-stretch interval and conditioning amplitude permutations were considered together (Fig. 4).

### Increased sampling density characterizes transitions from high to low muscle fiber resistance to stretch

In the data set above, we lacked data points between amplitude eliciting significant effects to characterize the transitions in the effect of conditioning stretch amplitude, motivating the additional data with finer sampling of conditioning stretch amplitude (Fig. 5).

**Figure 5:**
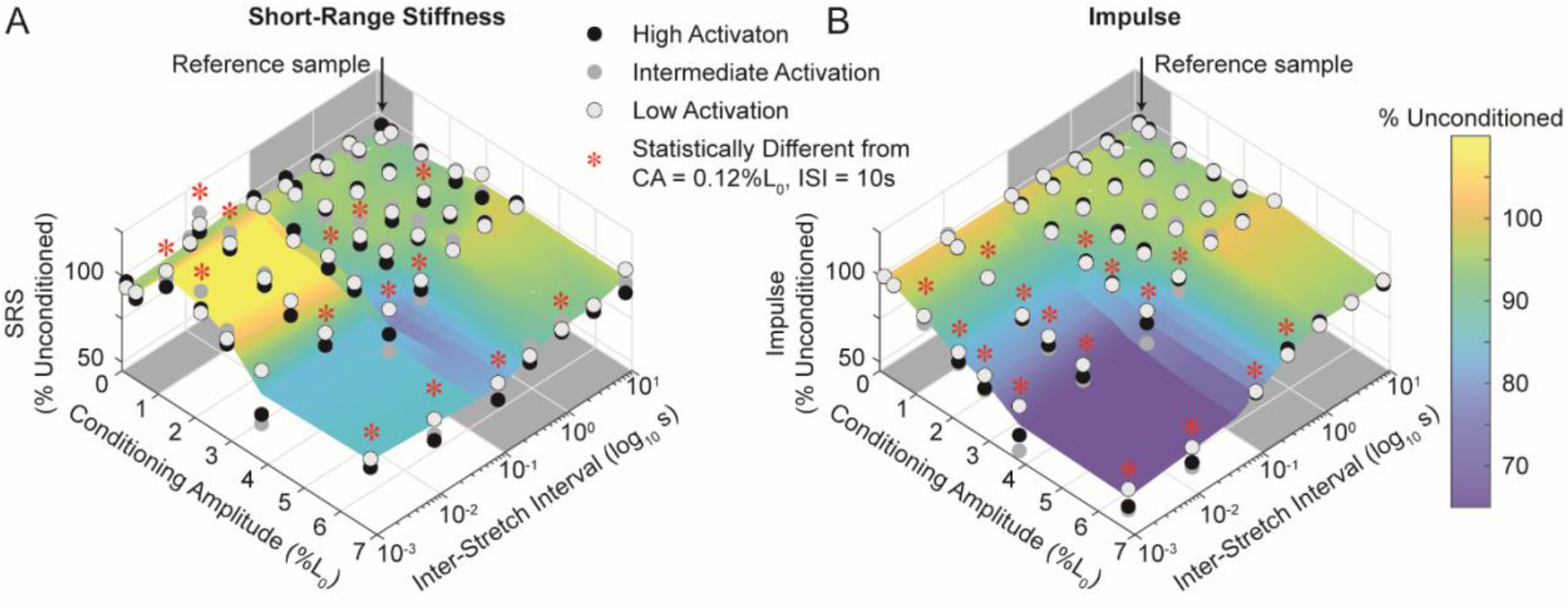
Increased sampling density reveals transitions in short-range stiffness and impulse with increasing conditioning stretch amplitude and decreasing inter-stretch interval. The effects of conditioning amplitude (left horizontal axis), inter-stretch interval (right horizontal axis) and activation level (colored circles; grey to black) on A) short-range stiffness and B) impulse as a percentage of unconditioned response (vertical axis). Gray boxes in both plots indicate posturally-relevant ranges of conditioning stretch amplitude (<1% L_0_) and inter-stretch interval (0.1 s to 1 s). Both SRS and impulse decreased monotonically as conditioning amplitude increased between 1-3% L_0_ but did not decrease further at 6% L_0_. This is similar to the nonlinear changes in stiffness as inter-stretch interval decreased, with no additional effects of decreasing the ISI below 0.01s.

Overall, the more densely sampled conditions in a subset of fibers (3 of 11) revealed a transition between high and low levels of relative resistance to stretch due to interactions between inter-stretch interval and conditioning stretch consistent with normal and abnormal postural sway and activation (Fig. 5). Specifically, the addition of more conditioning amplitudes between 1% and 7% L_0_ and a 0.01 s inter-stretch interval revealed a monotonic transition between high and low %unconditioned short-range stiffness (Fig. 5A) and impulse (Fig 5B), paralleling the reductions seen with varying inter-stretch interval (Fig. 5A, Fig. 5B, compare left and right horizontal axes).

Accordingly, the %unconditioned short-range stiffness was dependent upon interactions between inter-stretch interval and conditioning amplitude (F_36,731_=11.6, p < 0.001), as well as main effects of inter-stretch interval (F_6,731_=36.7, p < 0.001) and conditioning amplitude (F_6,731_=44.1, p < 0.001). Similar main and interacting effects of inter-stretch interval and conditioning amplitude were found for impulse (interaction: F_36,731_=20.9, p < 0.001; inter-stretch interval: F_6,731_=111, p < 0.001; conditioning amplitude: F_6,731_=115.8, p < 0.001);

Within the healthy posturally-relevant range where inter-stretch intervals were greater than 0.1 s and conditioning amplitudes were lower than 1% L_0_, both conditioned short-range stiffness (Fig. 5A) and impulse (Fig. 5B) remained high, near 100% unconditioned (Fig. 5, gray shaded regions). None of these conditions were significantly less than the reference condition of 0.001 s and 0.12% L_0_ conditioning stretch. Short-range stiffness was statistically higher than in the reference condition at the 0.01 s interval, 0.38% L_0_ amplitude permutation, which is within the posturally-relevant range.

The transition from high to low % unconditioned short-range stiffness and impulse occurred at combinations of shorter inter-stretch intervals and higher conditioning amplitudes characteristic of abnormal postural sway. Further, there appears to be a plateau in the reduction of muscle short-range stiffness at combinations of conditioning amplitude and ISI greater than 3% and lower than 0.01 s. The monotonic decrease and plateau effects could be readily observed in short-range stiffness across the 0.1s inter-stretch interval. Short-range stiffness was significantly reduced below 0.12%L_0_ levels at all conditioning amplitudes greater to or equal to 2.88%L_0_ (all p’s <0.05) when interval was fixed at 0.1 s. Short-range stiffness was also lower in the 6.82%L_0_ condition than at either 2.16% or 2.88%L_0_, demonstrating a persistent decrease in stiffness with increasing conditioning amplitude (Fig. 5A). However, there was no difference between short-range stiffness levels between the 3.83% and 6.82%L_0_ amplitudes, suggesting the effect reached a plateau (Fig. 5A, bottom). Impulse followed a similar pattern of monotonic decreases with increasing conditioning amplitude or decreasing inter-stretch interval, although no plateau effect was observed for impulse (Fig. 5B).

Finally, on average, relative change in short-range stiffness was highest in this sub-set of fibers at intermediate levels of activation when all conditioning amplitude and interval permutations were considered (F_2,732.4_=45.0, p < 0.001); with a similar effct of activation on impulse (Fig. 5B; STATS).

## Discussion

Here, we showed that history-dependant reductions in muscle resistance to stretch in single muscle fibers are absent in posturally-relevant activation and movement conditions but present in conditions consistent with abnormally elevated postural sway. We used a classical paired muscle stretch paradigm where a “conditioning” triangular stretch-shorten cycle is followed by a “test” ramp-and-hold. We systematically varied the triangular conditioning stretch amplitudes and inter-stretch intervals based on muscle stretch amplitudes and frequencies observed in normal and abnormal postural sway. The effects on short-range stiffness and impulse during the ramp-and-hold test stretch provided insight about how postural sway modulates the muscle resistance to stretch in a balance perturbation.

Our data revealed history dependent changes in muscle resistance to stretch when conditioning stretches occurred at larger and faster amplitudes than that seen in normal postural sway. Consistent with fluctuations in muscle length during standing postural sway in healthy individuals (Warnica et al., 2014), muscle short-range stiffness and impulse remained near 100% when the conditioning stretches were below ~1% fiber length and/or occurred greater than ~1 s prior to the test stretch at low, posturally-relevant activation levels (<15%). These findings were also consistent across all muscle activation levels. However, history-dependent reductions in muscle short-range stiffness and impulse were progressively observed as the conditioning stretch amplitude increased and/or inter-stretch intervals decreased. A maximum reduction to about 70% of nominal levels of short-range stiffness and impulse were observed when conditioning stretches were greater than 3% and occurred earlier than 0.1 s. Those with balance impairments are estimated to have ~2% fluctuations in muscle stretch during postural sway and higher sway frequency(Horak and Macpherson, 2011; Maurer et al., 2003), where muscle resistance reduced the most by pre-stretch conditions in our data. Notably, individuals with poor balance often increase their baseline muscle activity. Our data show that intermediate levels of activation (<40% max) increased absolute muscle resistance to stretch, but with greater history-dependent reductions in muscle resistance to stretch.

To our knowledge, this is the first study to characterize history-dependence of muscle resistance to stretch across multiple conditioning amplitudes, and their interaction with inter-stretch intervals. While prior studies demonstrated a minimum conditioning amplitude for inducing history-dependence in muscle fibers (Lakie and Robson, 1988), we revealed a transition region where muscle history-dependence was modulated by interactions between conditioning amplitude and inter-stretch interval. Within the set of experimental conditions tested, a variety of conditioning amplitude and inter-stretch interval combinations could cause changes in muscle resistance to stretch between 70-100% of unconditioned levels. In general, smaller stretches applied at long intervals preserved muscle resistance to stretch, whereas combinations of larger stretch with shorter inter-stretch intervals reduced muscle resistance to stretch. We did not test different stretch velocities, but based on crossbridge mechanisms (see below) slower stretches should tend to preserve muscle resistance to stretch, and faster stretches would tend to reduce muscle resistance to stretch (Rassier et al., 2003). Although we did not use continuous stretch trajectories, these data suggest that complex movement history of stretched muscle fibers such as that seen in postural behaviors will modulate muscle resistance to stretch.

To increase the behavioral relevance of our findings, we used two metrics of muscle resistance to stretch: short-range stiffness and impulse. Effects of muscle history-dependence to stretch have typically been studied using short-range stiffness or peak force (Campbell and Moss, 2002). Since muscle stiffness changes during stretch, neither of these metrics represent the overall force due to a stretch. Behaviourally, the success of a muscle in rejecting a postural perturbation depends on the overall effect due to stretch throughout a perturbation. The time-integral of muscle force i.e., impulse calculates the total force during the stretch and provides two insights relevant for sensorimotor control. It directly measures the change in momentum that the force can bring about on the body that it acts upon. Though not true in general, for our specific conditions where we use the same test stretch amplitude and velocity, differences in the muscle’s impulse and total work (calculated as force over distance) across conditions are proportional. Such impulse and work-based metrics have been critical is assessing muscle function in movement (Daley and Biewener, 2006; Dickinson et al., 2000).

Our results are consistent with predictions based on disrupting muscle cross-bridges. Briefly, muscle consists of elastic cross-bridges where myosin heads attach to actin sites to produce force. This force is assumed to be a function of the number and length of each attached cross-bridge. Muscle resistance to stretch arises when the attached cross-bridges at the onset of stretch are pulled to longer lengths and/or more cross-bridges become attached. However, sufficiently large amplitude stretch-shorten cause myosin heads to be unbound from actin, reducing muscle resistance to stretch. But, as inter-stretch interval increases, the cross-bridges begin to reattach, and over time, restores the muscle’s resistance to stretch to that of an unconditioned stretch. Since myosin heads must attach to actin to form cross-bridges, activation level can also affect the history-dependence of the muscle fiber. The number of available actin sites also affects both muscle force and the rate of recovery to steady-state length. At very low activation, there are not enough attached cross-bridges for the amplitude and inter-stretch interval conditionings to have much effect (Fig. 2C). At very high activation levels, there may be an abundance of actin sites available for cross-bridges to attach. This allows the cross-bridges to re-attach, reducing the effect of the conditioning stretch that disrupts the number and lengths of attached cross-bridges (Fig. 2C). Accordingly, we found a U-shaped relationship between muscle activation level and the history-dependence of muscle’s resistance to stretch. A more detailed mechanistic explanation of the biophysical mechanisms underlying history-dependent changes in muscle force can be found in a recent review (Lakie and Campbell, 2019).

Although our manipulations of muscle activation and conditioning stretches were based on the literature in postural control, there are a number of limitations when using isolated, permeabilized rat muscle fibers to infer the role of muscle properties on behavior. The forces acting on a human muscle-tendon unit during standing, as well as history-dependent changes to muscle resistance, depend on a range of other soft tissue properties, including muscle and tendon architecture, mechanical contributions from extracellular matrix and non-contractile proteins, as well as the muscle fiber type. Likewise, the continuous low and uniform chemical activation used in this study allowed us to manipulate the level of activation in fine detail, but it is unclear how the intermittent and unequal pulsatile activation seen across a whole muscle in behavior would affect muscle resistance to stretch. Furthermore, slow fibers used in standing balance could be active closer to 50% but would form a small percentage of all muscle fibers, making our findings here still relevant for postural control. Single muscle fibers data were also recorded at a relatively low temperature (22° C) to preserve fiber integrity; while there may be quantitative differences at body temperature, we expect the relative effects on conditioned versus unconditioned muscle resistance to stretch to be similar. Finally, holding muscle fibers isometric prior to, and between, stretches, is an artificial state that rarely occurs during behavior; we did not explicitly test the effects of continuous muscle stretching and shortening on muscle resistance to stretch.

Despite these limitations, our results may nonetheless offer some mechanistic insight into the role of pre-movement on the modulation of muscle properties in the context of balance control as well as other postural behaviors including those in the upper limb (Hu et al., 2011; Mathew and Crevecoeur, 2021). First, our data suggest that normal postural sway or movement variability is likely small enough that it does not induce history-dependent reductions in muscle resistance to stretch. Therefore, the mechanical resistance that muscle can provide is a critical first line of defense to stabilize the body before the slower, neurally-mediated balance corrections, even at low muscle activation levels (Ivanenko and Gurfinkel, 2018; Ting et al., 2009). Accordingly, recent work has shown a rapid increase in joint torque during balance perturbations prior to changes in muscle activation (De Groote et al., 2017; Van Wouwe et al., 2021). Second, our data suggest that increased postural sway consistent with that seen in balance impaired individuals reduced muscle resistance to stretch by up to 30%. Therefore, less mechanical stabilization of balance would be available for those with abnormally high postural sway. Accordingly, recent evidence shows that large, imposed, sway-like movements decrease ankle stiffness when people stand on a tilting platform (Sakanaka et al., 2016). Third, our data suggest that increased muscle activation to increase postural stiffness is more effective in healthy versus balance impaired individuals. People tend to increase muscle activity to decrease postural sway in threatening contexts such as standing at heights (Carpenter et al., 2001), when balance is evaluated by a clinician (Geh et al., 2011), or in response to arousing stimuli (Horslen and Carpenter, 2011). This increase in “postural stiffness” could be mediated by absolute changes in muscle resistance to stretch in conditions similar to normal postural sway (<1% L_0_, >1 s ISI). However, increased muscle activity observed in balance impaired individuals, may lead to a smaller decrease in postural sway because of the greater history-dependent effects of muscle resistance to stretch at conditions similar to abnormal postural sway (>3%, <1s ISI) at intermediate muscle activation levels. Overall, our data suggest that conditions of increased postural sway and muscle activation seen in balance impairments decrease the relative contributions of mechanical stabilizing mechanisms from muscle. As the history-dependent properties of muscle also contribute to the generation of sensory signals (Blum et al., 2020, 2017; Proske et al., 1993) that mediate balance-correcting responses (Lockhart and Ting, 2007; Welch and Ting, 2009), the neural response to balance perturbation is likely also diminished under conditions of increased postural sway. As such, the history-dependent properties of muscle resistance to stretch at elevated postural sway levels could negatively impact both mechanical and neural contributions to balance control.

## Abbreviations

SRS: Short-range stiffness
ISI: Inter-stretch Interval
SE: Standard Error
L_0_: Optimal Fiber Length
pCa: Potential of Calcium
SEM: standard error of the mean

## Acknowledgements

We would like to thank Friedl de Groote for her input to the manuscript and Faruk Moonschi for assistance with data collection.

